# Cheaters among pollinators: Floral traits drive spatiotemporal variation in nectar robbing and thieving in Afrotropical rainforests

**DOI:** 10.1101/2022.05.16.492032

**Authors:** Sailee P. Sakhalkar, Štěpán Janeček, Yannick Klomberg, Jan E.J. Mertens, Jiří Hodeček, Robert Tropek

## Abstract

- Nectar robbing and thieving can substantially affect the reproduction of animal-pollinated plants. Although the intensity of nectar exploitation remains unexplored at the community level, it probably varies along environmental gradients.
- We video-recorded flower visits to animal-pollinated plants in Afrotropical rainforests along a complete elevational gradient in the wet and dry seasons on Mount Cameroon. We analysed how the proportion of nectar robbing and thieving in the communities changes spatiotemporally, especially in association with the floral traits of the flowering plants.
- We recorded 14,391 flower visits, of which ~4.3% were from robbers (mostly bees and birds), and ~2.1% were from thieves (mostly flies, bees, and moths). Of the 194 studied plants, only 29 and 39 were nectar robbed and thieved, respectively. Robbers and thieves were most frequent at mid-elevations, with more frequent robbing in the wet season and thieving in the dry season. These trends were linked to the local composition of floral traits, and cheating groups’ associations to particular traits. Floral traits that prevented thieving made flowers susceptible to robbing, and vice versa.
- Spatiotemporal variation in floral traits across drives the cheating behaviour of flower visitors across communities, while indicating a trade-off between preventing nectar robbing and thieving.

## INTRODUCTION

Animal-pollinated plants use combinations of floral traits to advertise floral rewards, such as nectar, to attract pollinators (Willmer, 2011). However, conspicuous advertisement of nectar can also attract non-pollinators (Irwin & Maloof, 2002; Irwin *et al*., 2010) which may even dominate among all visitors (e.g., up to 78% of visitor species in Popic *et al*., 2013; King *et al*., 2013). These non-pollinators, called *cheaters*, do not touch floral reproductive organs while foraging on nectar (Irwin & Maloof, 2002; Irwin *et al*., 2010). Depending on how they extract nectar, such cheaters can be broadly categorised into either *robbers* or *thieves* (Inouye, 1980).

Thieves are visitors that can feed on nectar through the corolla opening without touching the stigmas or anthers while foraging because they are morphologically mismatched with the flowers they visit. Some mismatches are because the corolla tube is shorter than the visitors’ proboscis, as seen with lepidopterans which often probe for nectar without pollinating the flower (e.g., the *Eurybia lycisa* butterfly on *Calathea crotalifera;* Bauder *et al*., 2011), although some specialised plants rely on pollination by moths and butterflies (e.g. Balducci *et al*., 2019; Mertens *et al*., 2020). A morphological mismatch can also lead to thieving when small visitors feed on open-shaped flowers with exposed stigmas and anthers. For example, small meliponine bees visiting large, open Melastomataceae flowers (Murphy & Breed, 2008), and honeybees (*Apis mellifera*) visiting *Hypoestes aristata* flowers with protruding stigmas (Padyšáková et al. 2013) feeding on nectar without transferring pollen.

Robbers access nectar concealed in tubes or spurs of morphologically specialised flowers through holes in the floral structures, either made by themselves or by previous visitors (Rojas-Nossa *et al*., 2016). Opening holes in the corolla or calyx, termed *primary nectar robbing*, requires strong mandibles (e.g. bumblebees, carpenter bees, and beetles; Inouye, 1983), beaks (e.g. sunbirds, hummingbirds, and flowerpiercers; Geerts & Pauw, 2009; Janeček *et al*., 2011), or teeth (e.g. squirrels and galagoes; Deng *et al*., 2015). Once open, these holes can facilitate *secondary nectar robbing* by enabling other visitors to use them to access nectar (Bronstein, 2001; Irwin *et al*., 2010). Especially for visitors that can access nectar without piercing the flower, secondary robbing can be more energy-efficient than foraging through the corolla opening (Lichtenberg *et al*., 2018).

Nectar robbers and thieves lower nectar available to pollinators and change their foraging behaviour (Irwin *et al*., 2010; Padyšáková *et al*., 2013). In particular, floral damage caused during nectar robbing, besides inducing secondary robbing, makes flowers less attractive to subsequent visitors (Castro *et al*., 2013). Pollinators can learn to avoid such exploited flowers, leading to lower visitation rates and smaller pollinator communities for these flowers (Varma *et al*., 2020). Changes in pollinator behaviour due to cheaters can affect plants differently, with some benefitting from increased outcrossing and others exhibiting reduced pollen deposition and seed sets (Irwin *et al*., 2010). Whether nectar exploitation has positive or negative effects on plant reproduction depends on the species of the exploiters, pollinators and plants involved in the interactions and the variety of floral resources available. However, the intensity of such effects depends on how much nectar is removed, and therefore, on the visitation rates of cheaters (Maloof & Inouye, 2000).

Differences in rates of robbing and thieving of particular flowering plant species can be related to their floral traits. Thieving is more frequent on open-shaped flowers whose morphology makes them highly susceptible to morphologically mismatched visitors. In contrast, rewards in flowers with longer nectar tubes (Lázaro *et al*., 2015), fused petals, and closed shapes such as trap flowers (Gómez, 2005) are mainly available to their specialised pollinators. While these traits reduce nectar thieving, they can be more susceptible to robbers that cannot access the hidden nectar legitimately (Sonne *et al*., 2016; Rojas-Nossa *et al*., 2016). Thus, there could be a potential trade-off between floral traits that exclude nectar thieves and those that exclude nectar robbers.

Thieves and robbers are known to occur in many flowering plant communities around the world (Irwin & Maloof, 2002), yet community-wide studies of cheating behaviour are exceptionally rare. Nonetheless, the few existing studies suggest that nectar exploitation varies at several levels across communities (Irwin & Maloof, 2002; Rojas-Nossa *et al*., 2016; Cuta-Pineda *et al*., 2021). Rojas-Nossa *et al*. (2016) found that the proportion of plant species exploited within a given community varied geographically (e.g., 51.9% in the Mediterranean, 16.6% in the Alps, 22.2% in the Antilles, and 66% in the Andes). They further noted that within the set of exploited plants in each community, the level of exploitation varied across species (e.g., robbing on all flowers of *Thibaudia grandiflora* versus less than 5% of *Gaultheria erecta* flowers in the Andean community). Interestingly, the level of nectar exploitation was associated with the plants’ floral traits, with these trait associations differing across communities, suggesting that the floral traits influencing visitor behaviour can vary across communities. However, such studies are scarce and geographically biased towards the New World, with the Afrotropics being particularly understudied, making it difficult to infer general trends.

Our understanding of floral trait-pollinator behaviour associations is also restricted because spatiotemporal variation in the visitation rates of cheaters remains largely unexplored. Price *et al*. (2005) found variation in the visitation rates of nectar robbers on *Ipomopsis aggregata* among sites (0 to 0.06 robbing visits/flower/hour), as well as years (0.01 to 0.06 robbing visits/flower/hour at the same site across seven years). While (Castro *et al*., 2013)) found higher visitation frequencies of robbers on *Polygala vayredae* (on average 12.14 robbing visits/flower/hour), these also fluctuated in space (2.16 to 169.01 visits/flower/hour among three sites) and time (from 2.16 to 4.28 visits/flower/hour at the same site across three years). Although these changes could be due to available resources, environmental factors such as temperature and precipitation, as well as floral traits (Navarro, 2000), this has not been evaluated. Variation in the visitation rates of nectar robbers has been noted at the plant species level (Irwin & Maloof, 2002; Cuevas & Rosas-Guerrero, 2016), but nectar thieving has gone woefully unnoticed. Nevertheless, these studies indicate that visitation frequencies of cheaters on exploited plant species vary across communities, as well as fluctuate within and between seasons and years.

Mainly, the spatiotemporal variation of nectar robbing and thieving has not been quantified at the community level, although it can be expected to follow spatiotemporal changes in floral traits of plant communities (Irwin & Maloof, 2002). For instance, the community composition of floral traits varies along elevational gradients (Albrecht *et al*., 2018; Klomberg *et al*., 2022). At higher elevations, lower temperatures raise the energetic requirements of pollinators (Classen *et al*., 2015). This may increase the importance of traits related to nectar production and composition for floral visitors (Klomberg *et al*., 2022). Floral traits can also vary seasonally, as some plants blooming in rainy seasons have closed flowers with narrow tubes (Klomberg *et al*., 2022) that prevent nectar dilution (Aizen, 2003) and washing away of pollen (Mao & Huang, 2009). In the tropics, ornithophilous plants are commoner in the wet season, as seen with sunbird-pollinated plants on Mount Cameroon (Janeček *et al*., 2021).

In this study, our main aim was to quantify nectar exploitation by cheaters in the Afrotropical rainforests along an elevational gradient on Mount Cameroon during dry and wet seasons. We addressed the following questions: (a) How frequent are nectar robbing and thieving in the studied communities? (b) Do the rates of nectar exploitation behaviours vary along the elevation or between seasonal extremities? (c) Is spatiotemporal variation in visitor behaviour associated with the floral traits of the studied plants? (d) Do nectar robbers and thieves differ in their associations with specific floral traits, and do these associations change with their functional groups?

## METHODS

### Study area and sites

We studied flower-visitor communities on Mount Cameroon (Southwestern Region, Cameroon; 4°12’10” N, 9°10’11” E), the highest mountain in West and Central Africa (4,095 m a.s.l.). The mountain’s southwestern slope is the only continuous elevational gradient in continental Africa, with pristine tropical rainforest extending from the lowland (~350 m a.s.l.) to the timberline (~2,100 to 2,300 m a.s.l.). Precipitation at our study area is strongly seasonal, with over 2,000 mm monthly rainfall in the foothills during the wet season (June to September) and little to none in the dry season (November to February;(Maicher *et al*., 2016, 2020).

Along the rainforest gradient, we sampled four elevations: lowland (650 m a.s.l), sub-montane (1,100 and 1,450 m a.s.l.), and montane (2,200 m a.s.l.), once each in the wet and dry seasons (for details, see Table S1, and Klomberg *et al*., 2022). At each elevation, we established six transects (200 × 10 m) spaced at least 100 m apart to represent the local vegetation heterogeneity.

### Behavioural observations

We collected data on flower-visitor interactions using security cameras (VIVOTEK IB8367RT with IR night vision) to video-record all zoophilous plant species in flower across all vegetation strata for continuous 24-hour periods. We set the video frames to record the maximum number of flowers allowing visitor identification from videos (for a detailed methodology, see Klomberg *et al*., 2022; Mertens *et al*., 2021). We recorded flowers along the established transects at each elevation, aiming to record five replicates per species per site and season, including the surrounding vegetation, when the number of replicates for the observed plant species was insufficient. Flower-visitor interactions were observed from the recordings with motion detection (Motion Meerkat 2.0; Weinstein, 2015) when possible, and manual playback when not.

We identified visitors to the best taxonomic resolution, sorting them into morphospecies when possible. We then sorted them into 13 functional groups following common pollination syndromes (Willmer, 2011), splitting bees and flies into subgroups that better represent the difference in their reward preferences: sunbirds, bats, small mammals, hoverflies, other flies (hereafter ‘flies’), honeybees, carpenter bees, other bees, beetles, wasps, butterflies, hawkmoths, and other moths (hereafter ‘moths’). To account for differences in the sampling effort among individual sites and seasons (different numbers of plant species in flower), and plant species (different numbers of recorded flowers, differences in flower longevity, technical failures, and/or lack of enough replicates for rare plants), we quantified the visitation frequency for each behaviour as the number of visits per flower per minute.

Based on their foraging behaviour, we sorted all floral visitors into three groups: (a) *pollinators* – contact with anthers and/or stigmas; (b) *thieves* – nectar accessed through the floral opening without touching anthers or stigmas; and (c) *robbers* – nectar accessed through holes other than the floral opening and without touching any reproductive organs (Fig 1 B-E). All other visitors that neither approached floral rewards nor touched reproductive organs were excluded. Further, we defined the *main pollinators* of each plant species as the two functional groups with the highest frequencies of pollinatoing visits.

**Figure 1.**
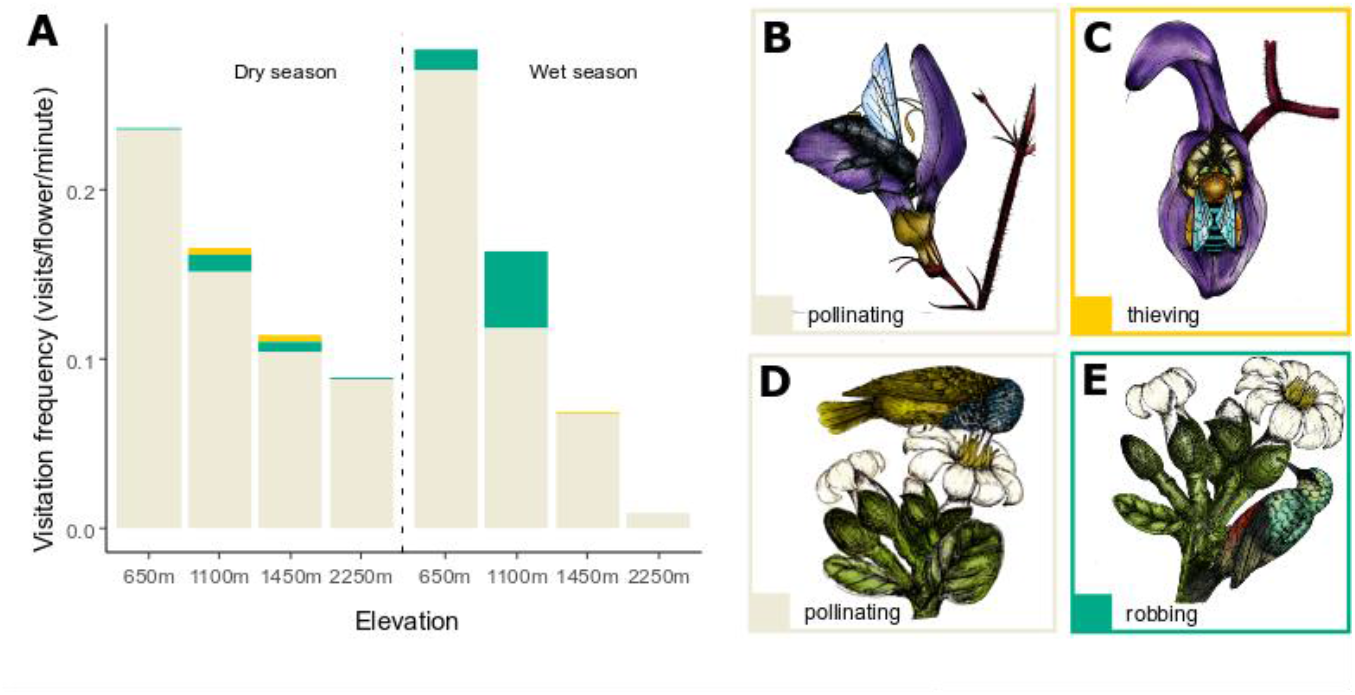
Robbers, thieves, and pollinators of flowering plants on Mount Cameroon. (A) Visitation frequencies for pollinators (grey), thieves (yellow), and robbers (blue) across the elevational gradient and between the dry and wet seasons on Mount Cameroon. Examples of flower visitors: (B) a carpenter bee (*Xylocopa* sp.) pollinating and (C) a blue-banded bee (*Amegilla* sp.) thieving nectar of *Brillantaisia owariensis*. (D) A Cameroon Sunbird (*Cyanomitra oritis*) pollinating and (E) a Northern Double-Collared Sunbird (*Cynniris reichenowi*) robbing nectar of *Anthocleista scandens* (all drawings made by Sailee P. Sakhalkar).

#### Floral traits

We measured ten floral traits (Table S2; partly used in Klomberg *et al*., 2022). Three of these were morphometric (corolla size, and the length and width of nectar tube), and seven were qualitative characterising floral shape (bell, bowl, dish, funnel, gullet, labiate, open, papilionaceous, salverform, stellate, trumpet, tube, urceolate), symmetry (zygo- and actinomorphy), orientation (horizontal, pendant, upright), colour (brown, green, orange, pink, purple, red, white, yellow), nectar guides (presence/absence), brightness (vivid/drab), and odour strength (none, weak, strong), and amount of nectar sugar (measured as 24-hour production in flowers in-situ, as described in (Janeček *et al*., 2021).

#### Data analyses

We tested if the relative proportions of visitor behaviour (robbing, thieving, and pollinating) differed spatiotemporally using Chi-square goodness-of-fit tests (the *chisq.test* function from the *stats* package). We separately tested for differences between the two seasons, and among the four elevations.

We analysed interspecific trait-behaviour associations using the multivariate RLQ analysis to test whether community-level differences in visitor behaviour are associated with floral traits (Dolédec *et al*., 1996). RLQ is an ordination method that associates matrices **R** and **Q** containing two separate sets of variables (such as environmental variables and species traits) by maximising their covariance based on a third, central matrix **L** containing variables (such as species composition) that link **R** and **Q**. In our study, matrix **R** (*n* × *m*), where for each of *n* sites, *m* columns contained data for the frequencies of total visitor behaviour (pollination, robbing and thieving) and environmental variables (season and elevation); **L** (*n* × *p*) contained presence-absence data each from *n* sites for *p* plant species; and **Q** (*p* × *q*) that had measurements of *p* plant species for *q* floral traits.

Firstly, we analysed plant species composition at each site (L) using correspondence analysis (CA). Secondly, we used the row weights (sites) and column weights (plant species) from the CA to weight the rows of matrices R (sites by environmental variables and visitor behaviour) and Q (plant species by floral traits). Thirdly, we used Hill-Smith analyses for the separate ordinations of R and Q as they contained both continuous and categorical variables. Next, in the final step of the RLQ analysis, these separate ordinations were combined by performing a double inertia analysis of R and Q, linked through L. The analysis found those linear combinations of the environmental variables and floral traits that maximise their covariance, thus describing their joint structure (overview in Fig.S1). We evaluated the significance of the RLQ analysis using a sequential two-model Monte Carlo test with 9999 permutations to test if species composition was linked to trait composition (by permuting species) as well as to environmental and behavioural variables (by permuting sites). Lastly, we used ordination diagrams to visually examine the joint structure of the three matrices (Fig. 2A), and the associations between sites and visitor behaviour (Fig. 2B) and between sites and floral traits (Fig. 2C). The RLQ analysis was conducted using the *ade4* package (Dray & Dufour, 2007).

**Figure 2:**
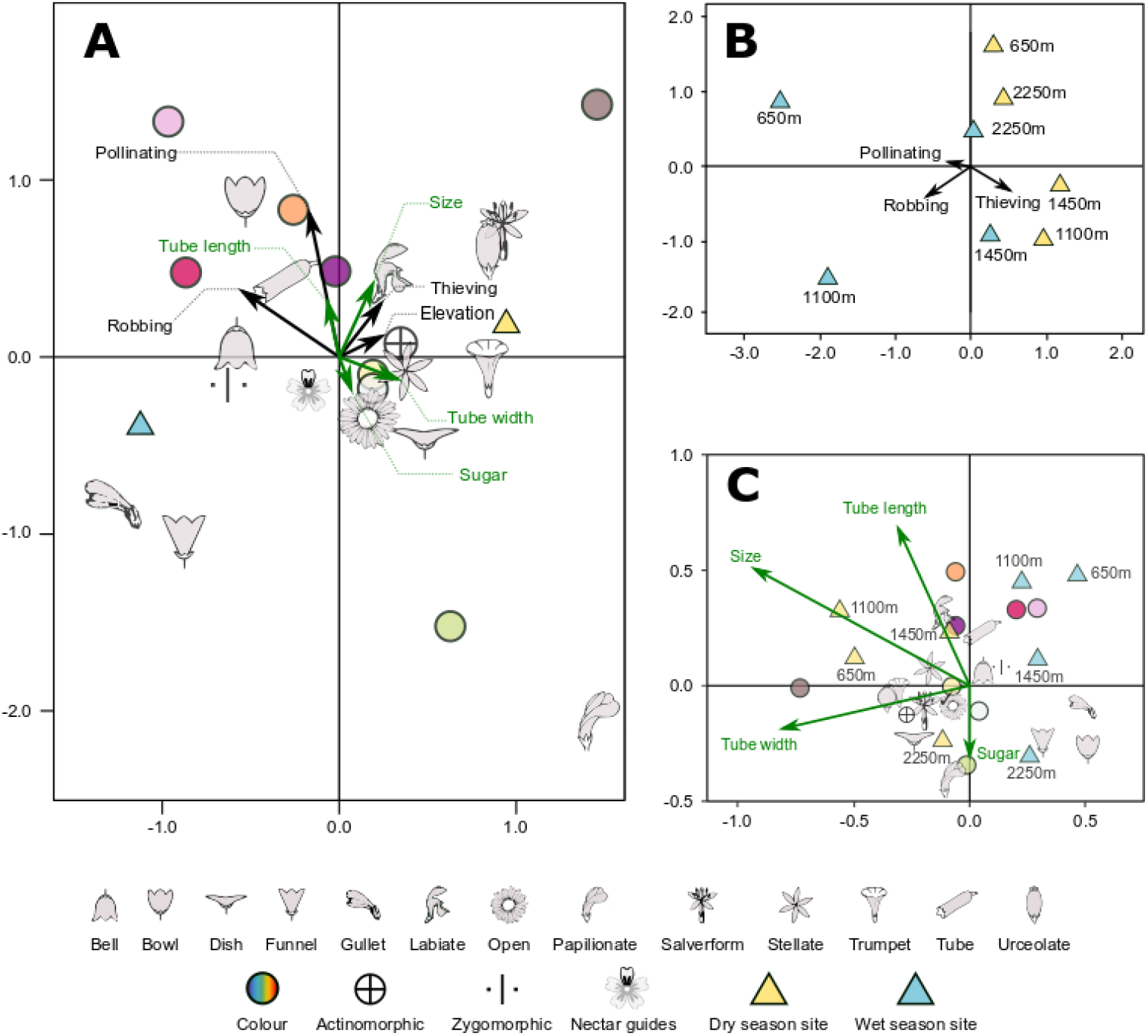
Ordination diagrams (biplots) visualising the associations between environmental variables, visitation frequencies and floral traits on Mount Cameroon, as resulting from the RLQ analysis. Season, categorical floral traits, and site centroids are represented by symbols as seen in the legend, whereas continuous floral traits (in green) and visitation frequencies (in black) are visualised with arrows. (A) Associations of environmental variables and visitation frequencies with floral traits. (B) Associations of sites with visitation frequencies and environmental variables. (C) Associations of sites and floral traits. Dashed lines are used to label the arrows to improve readability.

Finally, we used separate Redundancy Analyses (RDA) for robbers and thieves to test whether each of their functional groups has different trait associations. In these independent analyses, robbing and thieving frequencies of functional groups per plant species were Hellinger-transformed (for zero-inflated data with low counts) and used as response variables. In both analyses, floral traits were used as explanatory variables and were chosen based on forward selection. We tested the significance of each RDA using Monte Carlo tests with 999 permutations. The RDAs were performed using the *vegan* package (Oksanen *et al*., 2019).

All statistical analyses were conducted in R 4.1.1 (R Core Team, 2021).

## RESULTS

We video-recorded flowers for a total of 26,138 hours (i.e., >2.98 years) along the elevational gradient of Mount Cameroon. These recordings resulted in a total of 14,391 visits of which 13,365 (92.87%) were pollinators, 623 (4.32%) robbers, and 304 (2.11%) thieves. Of a total of 195 plants, only 26 (14.79%) were robbed, while 39 (19.89%) were thieved, while neither thieves nor robbers were observed for 126 plants (64.28%) (Table 1). There were also 12 plants that were not visited by robbers, thieves, or pollinators. On average, 13% of visitors to exploited plants were thieves, while 24% were robbers, but the proportions of exploiting visitors differed among plant species. Some key examples of visitors can be found in our video (Sakhalkar *et al*., 2022). While *Plectranthus decurrens* had the highest proportion of visits (over 90%) from robbers, *Crassocephalum montuosum* had the lowest (0.12%). Similarly, *Pararistolochia zenkeri* was visited only by nectar thieves, while only 0.03% of visits to *Psydrax dunlapii* were from thieves.

**Table 1:** Overview of flower visitors on Mount Cameroon. Visitation frequencies (i.e. number of visits per flower per minute) and numbers of visits, numbers of visiting morphospecies, and numbers of visited plants are listed separately for pollinators, robbers, and thieves in each functional group of flower-visitors.

| Taxonomic group | Functional group | Visitation frequency (number of visits) |  |  |  | Visitor morphospecies |  |  |  | Visited plant species |  |  |  |
| --- | --- | --- | --- | --- | --- | --- | --- | --- | --- | --- | --- | --- | --- |
|  |  | Pollinator<br>s | Robber<br>s | Thieve<br>s | Total | Pollinator<br>s | Robber<br>s | Thieve<br>s | Total | Pollinator<br>s | Robber<br>s | Thieve<br>s | Total |
| **Coleoptera** | Beetles | 0.0234<br>(267) | - | 0.0036<br>(33) | 0.027<br>(300) | 38 | 0 | 7 | 70 | 19 | 0 | 7 | 48 |
| **Chiroptera** | Bats | 0.0033<br>(39) | - | - | 0.0033<br>(39) | 1 | 0 | 0 | 1 | 1 | 0 | 0 | 1 |
| **Diptera** | Flies | 0.1251<br>(1308) | - | 0.0073<br>(62) | 0.1324<br>(1370) | 29 | 2 | 9 | 31 | 62 | 2 | 18 | 96 |
|  | Hoverflies | 0.3077<br>(2097) | 0.0001<br>(1) | 0.0077<br>(61) | 0.3155<br>(2159) | 66 | 0 | 16 | 88 | 41 | 0 | 20 | 88 |
| **Hymenoptera** | Carpenter bees | 0.0325<br>(155) | - | - | 0.0325<br>(155) | 0 | 0 | 2 | 2 | 9 | 0 | 0 | 20 |
|  | Honeybees | 0.2398<br>(3397) | 0.0071<br>(123) | 0.0034<br>(58) | 0.2503<br>(3578) | 1 | 1 | 1 | 1 | 27 | 3 | 1 | 47 |
|  | Other bees | 0.5461<br>(3402) | 0.0565<br>(400) | 0.0028<br>(43) | 0.6054<br>(3845) | 19 | 6 | 4 | 19 | 73 | 15 | 2 | 100 |
|  | Wasps | 0.0711<br>(450) | 0.0032<br>(24) | - | 0.0743<br>(474) | 19 | 4 | 0 | 21 | 12 | 4 | 0 | 30 |
| **Lepidoptera** | Butterflies | 0.0713<br>(595) | - | 0.0011<br>(17) | 0.0724<br>(612) | 77 | 0 | 8 | 80 | 22 | 0 | 3 | 59 |
|  | Hawkmoths | 0.0113<br>(6) | - | - | 0.0113<br>(6) | 26 | 0 | 0 | 26 | 31 | 0 | 0 | 31 |
|  | Moths | 0.0731<br>(1408) | 0.0004<br>(5) | 0.0018<br>(30) | 0.0753<br>(1443) | 257 | 5 | 19 | 283 | 39 | 3 | 6 | 80 |
| **Passeriformes** | Sunbirds | 0.0357<br>(229) | 0.0075<br>(58) | - | 0.0432<br>(287) | 4 | 5 | 0 | 5 | 12 | 7 | 0 | 23 |
| **Other mammals** | Small mammals | 0.0024<br>(12) | 0.0014<br>(12) | - | 0.0038<br>(24) | 3 | 4 | 0 | 8 | 3 | 2 | 0 | 6 |
| **Total** |  | 1.543<br>(13464) | 0.0761<br>(623) | 0.0278<br>(304) | 1.647<br>(14391)<br>) | 540 | 27 | 66 | 635 | 180 | 29 | 39 | 195(of<br>which 12<br>not visited<br>by robbers,<br>thieves or<br>pollinators.<br>) |

All 19 recorded morphospecies of other bees were pollinators of at least part of the visited plants. This functional group was the main (i.e., the first or second most frequent) pollinator for 73 plant species in our study and the most frequent pollinators (details on all visiting groups are summarised in Table 1). The second most frequent group of pollinators was hoverflies, with nearly all morphospecies pollinating flowers and 62 plant species relying on them as their main pollinators. Although the most pollinator morphospecies were moths, they were the main pollinators of only 39 plant species. Interestingly, other bees were also the most frequent robbers, while hoverflies were the most frequent thieves. While sunbirds and honeybees were equally frequent robbers, honeybees robbed only three of the 47 plant species they visited, while sunbirds robbed seven of 23 visited plant species. Although nectar robbing (0.0761 visits/flower/min) was four times more frequent than thieving (0.0278 visits/flower/min), 66 morphospecies thieved flowers as compared to 27 morphospecies that robbed. After hoverflies, the most common thieves were flies, honeybees, and beetles. While 19 morphospecies of moths and eight butterflies thieved flowers, they were the least frequent nectar thieves (Table 1).

### Spatiotemporal variation in visitor behaviour

Total visitation frequency and pollination frequency declined with increasing elevation and were lower in the dry season than in the wet season (Fig. 1A, Table 2). However, this pattern differed for robbing and thieving. In the dry and wet seasons, robbing frequency increased from 650 m to 1,100 m a.s.l. and then declined to 2,250 m a.s.l. Robbing frequency was also higher in the wet season compared to the dry season. On the other hand, thieving frequency peaked at 1,450m in both seasons and was more frequent in the dry season (Fig. 1A, Table 2). These spatiotemporal differences were also statistically significant for the relative ratios of visitor behaviour between the seasons (chi-square = 474.63, *df* = 3, p-value < 0.001) as well as among the elevations (chi-square = 2946.3, *df* = 9, p-value < 0.001).

**Table 2.**
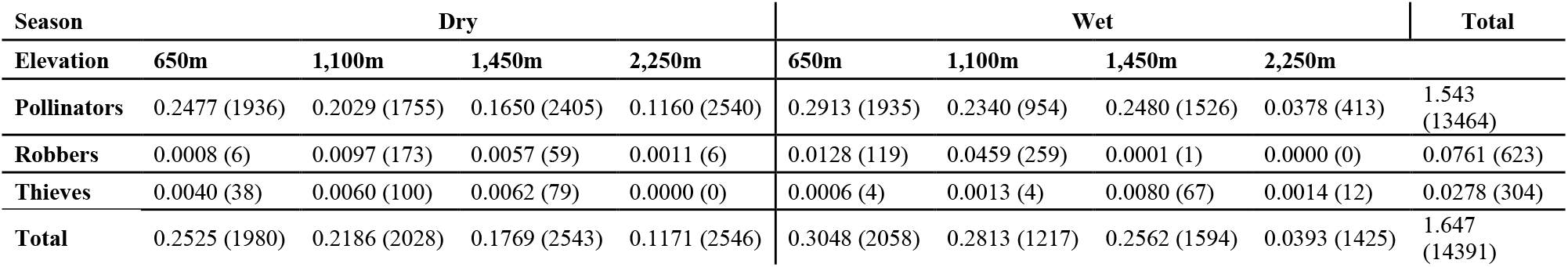
Visitation frequencies (i.e. number of visits per flower per minute) and number of visits (in brackets) for pollinators, robbers, and thieves at each of the four elevations and two seasons sampled on Mount Cameroon.

### Association of visitor behaviour, floral traits, and spatiotemporal variation

The associations between environmental variables (elevation and season) and visitor behaviour (*F* = 0.705, *p* = 0.042), and between environmental variables and floral traits (*F* = 0.705, *p* = 0.001) were statistically significant. As summarised in Table S3, the first RLQ axis explained 96.67% and 62.92% of the first-axis variation from the separate analyses of the variation in environmental variables and visitor behaviour (Fig. 2B), and in floral traits (Fig. 2C), respectively. The two RLQ axes explained 81.22% (Table S3; Fig. 2A) of the joint structure, with the first axis capturing most of the variation. This showed that their joint structure was strongly associated through the plant species composition across the sites.

The ordination diagrams (Fig. 2) show that the first axis separated communities according to the two seasons (Fig. 2B). Nectar robbing was associated with the wet season and flowers that were zygomorphic, bell-shaped, or tubular, and orange or red (Fig. 2A), and increased with an increase in tube length. Nectar thieving was closely associated with trumpet-shaped, urceolate, and open flowers and appeared to be influenced by flower size and tube width. While pollination was more frequent in the wet season (Fig. 2B), it had a closer association to actinomorphic flowers that were salverform, stellate, or dish-shaped (Fig. 2A). There was no apparent elevational pattern in the association of floral traits and visitor behaviour.

### Floral trait associations of floral cheaters

Robbers had clearer associations to plant species with specific pollinating groups than thieves (Fig 3A, 3C), although this pattern varied among the functional groups of cheaters. As nectar robbers, moths exploited plant species mainly pollinated by honeybees, other bees, butterflies, and sunbirds. However, moths were less selective while thieving, exploiting plants with main pollinators from all functional groups except hawkmoths, wasps, bats, and small mammals. This was also the trend for hoverflies, as they robbed plant species with either flies, hoverflies, or sunbirds as their main pollinators. At the same time, they thieved plants that were mainly pollinated by flies, butterflies, moths, carpenter bees, honeybees, and small mammals. Unlike hoverflies and moths, other bees were more selective while thieving plants rather than robbing them. Other bees were robbers of plants that all functional groups besides flies, small mammals, and honeybees pollinated. However, other bees only thieved plants mainly pollinated by themselves, carpenter bees, and hoverflies.

**Figure 3:**
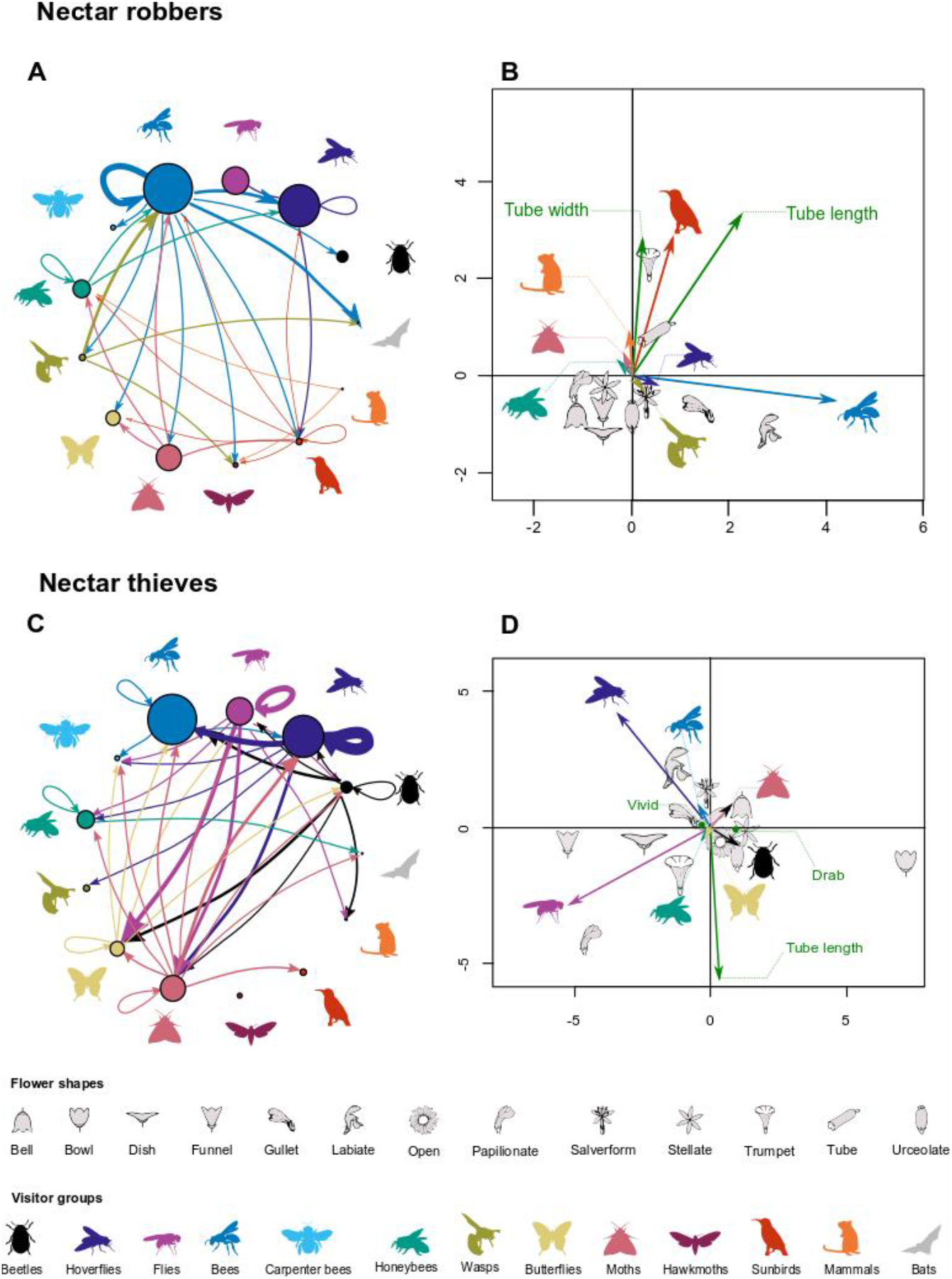
Floral preferences of nectar robbers and nectar thieves based on the main pollinators of the exploited plants (circular graphs), and trait associations for functional groups of robbers and thieves (ordination diagrams). A, C: Robbers are more specific than thieves with their choice of exploited plant species. The circular graphs visualise how functional groups of cheaters differ in their choice of robbing (A) or thieving (C) plants with main pollinators from specific functional groups. All functional groups are visualised as silhouettes, with circles representing the number of plant species for which each group is a main pollinator. Arrows connect to circles containing the plant species that a functional group of robber or thief exploits; their thickness corresponds to the number of exploited plant species. (B, D): Ordination diagrams (RDA) visualising the association between floral traits and the frequency of exploitation for different functional groups of robbers (B) and thieves (D). Arrows visualise the frequencies of robbing or thieving for of each functional group (coloured as silhouettes) and continuous floral traits (in green). Dashed lines are used to label the arrows to improve readability. The centroids for flower shapes, and the labels for functional groups are marked with symbols as shown in the figure legend.

Floral traits were significantly associated to the frequencies of nectar robbing and thieving by different functional groups of visitors (RDA for robbing: F = 3.22, *p* = 0.001, 16.56% of explained variation; RDA for thieving: F = 2.30, *p* = 0.001, 12.00% of explained variation; Table S4). However, these associations were clearer for some groups of cheaters than others. Functional groups of nectar robbers had different associations to flower shape, tube length, and tube width, with the relationship being clearer for other bees and sunbirds (Fig. 3B). Sunbirds robbed mainly tubular flowers and those with wide and long nectar tubes. Other bees did not have a strong association to robbing flowers with larger nectar tubes and appeared to rob gullet-shaped and labiate flowers. Although nectar thieving groups had different associations with flower brightness, shape, and tube length (Table S4, Fig. 3D), these were strongest for flies and hoverflies. In general, flowers with longer tubes were not associated with thieving by any of the functional groups, with floral shape influencing which functional group would thieve the flower. Flies thieved funnel-shaped and papilionate flowers, hoverflies were thieves of labiate flowers, and other bees thieved gullet and salverform flowers. Drabness separated flowers that beetles thieved from those thieved by other groups. The traits associated with thieving by butterflies and moths were not apparent from the ordination diagram (Fig 3D).

## DISCUSSION

We revealed that nectar robbing and thieving were rare in tropical communities on Mount Cameroon. Moreover, we found that their frequency varied spatiotemporally, probably driven by the floral trait distribution in communities which affected the behaviour of nectar foragers.

### The rarity of nectar exploitation in communities

We observed nectar robbing and thieving across all communities, regardless of the season and elevation (Fig. 1 A). Our observation is consistent with previous findings which suggested that cheaters probably occur in all flower-visitor communities (Irwin *et al*., 2010; Rojas-Nossa *et al*., 2016). However, in our study, most plant species (64%) were visited only by their potential pollinators. The frequencies of robbers and thieves were consistently low even on plant species that were exploited by cheaters. Although cheaters outnumbered pollinators for some previously studied plant species (Castro *et al*., 2013; Cuevas & Rosas-Guerrero, 2016), only a subset of plants were exploited, similar to our results. Although Rojas-Nossa *et al*. (2016) found 16-66% of plant species robbed across communities, they only included species morphologically vulnerable to nectar robbing. Thus, the proportion of all robbed plant species is most probably much smaller, similar to our findings (i.e. 14.79%).

### Variation in cheating frequencies across space and time

The frequencies of cheaters substantially differed between the seasons, with a less clear elevational trend. Nectar robbing was higher in the wet season, whilst the opposite pattern was observed for thieving. We found that the spatiotemporal differences in the floral visitors’ behaviour can be related to the spatiotemporal distribution of floral traits in the plant communities. Thus, our results support suggestions from the previous synthesis on an inter-biome scale (Rojas-Nossa *et al*., 2016) that the key determinants of nectar robbing among communities could be morphological adaptations of flowers for their specialised pollinators. On Mount Cameroon, morphologically generalised flowers prevail in the dry season, whilst specialised flowers in the wet season. Floral specialisation helps reduce the number of thieves and ineffective pollinators (Lázaro *et al*., 2013), but the unexploited nectar can attract robbers (Irwin *et al*., 2010).

In our study, nectar robbing was more frequent in the wet season, where flowers with longer nectar tubes and generally narrower corollas were more common (Fig 2A). Such flowers were prevalent in the wet season, where their morphology could help avoid nectar dilution (Aizen, 2003) or specialise to their pollinators with higher energetic needs (Chmel *et al*., 2021). However, such specialised flowers are more often robbed by small-tongued visitors unable to reach their nectar through the flower opening (Bronstein, 2001; Navarro & Medel, 2009; Maruyama *et al*., 2015). We found a weak association between robbing and flower size. Larger flowers are expected to produce more nectar (Harder & Cruzan, 1990), which could compensate for the effort of making a hole in the flower to reach the nectar, making robbing profitable for visitors such as bees (Irwin *et al*., 2010). Indeed, we found that some larger flowers such as *Impatiens* spp. with narrow and curved spurs (for sunbirds;(Janeček *et al*., 2015), or *Kigelia africana* with relatively easily accessible nectar within the large floral chamber (evolved for bats; Newman *et al*., 2021), were robbed (see video in Sakhalkar *et al*., 2022).

We found nectar thieving more common in flowers with open and trumpet shapes, larger sizes, and wider tubes, which prevailed in the dry season (Fig. 2B). Such flowers are prone to morphologically mismatched nectar thieves (Irwin *et al*., 2001). In our study, larger, nectar of open flowers with reproductive organs clustered at the flower centre (e.g., *Oncoba lophocarpa*) was thieved relatively often. Another notable mismatch due to the small size of visitors was observed in large and trumpet-shaped flowers of *Kigelia africana* commonly visited by relatively smaller honeybees without touching the long, filamentous anthers. Further, nectar of gullet flowers of *Brillantaisia owariensis* is often thieved by small pollinators, such as blue-banded bees (*Amegilla* spp.) and skippers (Lepidoptera: Hesperiinae), too small to touch reproductive organs of this plant usually pollinated by large carpenter bees (*Xylocopa* spp.) (Fig. 1 B-C; see video in Sakhalkar *et al*., 2022).

### Floral preferences of functional groups of cheaters

We expected smaller bees (such as halictids) to thieve open flowers successfully. However, such bees only robbed two plant species (*Brilllantaisia owariensis* and *Plectranthus kamerunensis*) where they do not touch the reproductive organs adapted for larger bees (see video in Sakhalkar *et al*., 2022). On the other hand, bees were the most frequent nectar robbers in our study, especially in flowers with longer tubes (e.g., *Bertiera racemosa*), and labiate and gullet shapes (e.g. *Impatiens niamniamensis, P. kamerunensis*). Although we did not directly test the effect of available nectar volume on bee behaviour, we observed halictid bees on the labiate flowers of *Plectranthus* spp. feeding through the flower opening, and switching to robbing when its nectar level became shorter and inaccessible (see video in Sakhalkar *et al*., 2022). Such behaviour switching by bees in relation to the resources available has already been described (Bronstein *et al*., 2017; Lichtenberg *et al*., 2020). All six morphospecies of robbing bees behaved also as potential pollinators in our study, suggesting that for some plant species, bees can be antagonists (robbers) as well as mutualists (pollinators). However, this cannot be further discussed without experimental data.

Sunbirds were the second most frequent robbers in our study, in concordance with other studies (Geerts & Pauw, 2009). Although flowers can often be adapted to ornithophily, sunbirds are attracted to flowers with a high amount of nectar, regardless of other floral traits ((Chmel *et al*., 2021b). Thus, when their bills are too short to access nectar within long-spurred flowers, they often resort to nectar robbing (Janeček *et al*., 2015). Interestingly, sunbirds also robbed larger flowers with accessible nectar, such as *Kigelia africana, Costus dubius*, and *Anthocleista scandens* (Fig. 1. D-E; see video in Sakhalkar *et al*., 2022). While such flowers usually allow sunbirds to feed legitimately and with relatively short handling times (Temeles & Pan, 2002), wider flowers might offer more nectar, making them more likely to rob.

While we studied some traits that made flowers susceptible to robbing, we did not measure various floral traits protecting against nectar robbing, such as petal thickness, calyx density, the packing of flowers in inflorescences, and corolla stickiness (Rojas-Nossa *et al*., 2016; McCarren *et al*., 2021). Such ‘protective traits’ could explain why only a fraction of flowers susceptible to robbing were exploited, although we could not evaluate their importance.

The most frequent nectar thieves in our study were hoverflies and flies, although we did not find any mention of flies as thieves in other studies. Typically, flower-visiting flies are attracted to open flowers with easily accessible nectar (Branquart & Hemptinne, 2000). The declining frequency of thieving by non-specialised flies and hoverflies with nectar tube length was predictable, considering that they would be unable to feed on nectar with small proboscides (Doyle *et al*., 2020). Despite this, longer flowers may still be thieved by small-sized flies and hoverflies if their nectar tubes (e.g. *Aframomum* spp.) or spurs (*Impatiens* spp.) are wide enough (Zhang *et al*., 2014), or if they produce enough nectar accumulate in the tube (Vlašánková et al., 2017). Similarly, small beetles were also common nectar thieves in generalised open flowers (e.g. *Begonia* spp.), as shown by numerous other studies (Gómez, 2005; Sayers *et al*., 2019). Additionally, we observed small beetles to also thieve nectar of some closed flowers with large chambers (e.g., *Aframomum* spp.). None of the beetles in our study robbed flowers, although a study reports nectar robbing by weevils (Clement, 1992). Surprisingly, we found that moths and butterflies were not very frequent nectar exploiters (Mertens *et al*., 2021). Although lepidopterans can be efficient pollinators (Schemske, 1976; de Araújo *et al*., 2014), many lepidopterans with long proboscis can feed on diverse flowers, generalised or specialised, without pollinating them (Bauder *et al*., 2011). All flowers thieved by butterflies and moths in our study were either large and trumpet-shaped (e.g., *Pararistolochia zenkeri*) or had exposed reproductive organs that allowed bypassing pollination (e.g., *Clematis simense*) and caused the morphological mismatch (Irwin *et al*., 2010).

### Conclusions

We found the nectar thieving and robbing generally uncommon in the Afrotropical forests. Nevertheless, the revealed spatiotemporal variation in visitor behaviour can arise from the uneven distribution of floral traits in communities. The higher prevalence of closed flowers in the wet season was associated with robbing, whilst more open flowers in the dry season was related to more common nectar thieves. This confirmed the trade-off between the floral traits that limit robbers and those that restrict thieves. We also related specific floral traits to the robbing and thieving behaviour of particular functional groups of nectar exploiters. Altogether, we demonstrated that floral traits that are commonly known to shape plant-pollinator interactions significantly influence nectar robbers and thieves.

## ACKNOWLEDGEMENTS

We thank Ishmeal N. Kobe, Nestoral T. Fominka, Vincent Maicher, Luma Francis Ewome, Raissa Dywou Kouede, Esembe Jacques Chi, Karolina Hrubá, Hernani Oliveira, Zuzana Sejfová, Pavel Potocký, Pavel Kratochvíl, and other assistants for their help in the field; Ivan Šonský, Petra Janečková, Eliška Chmelová, Marek Rybár, Jan Raška, and several others for their assistance in processing video recordings; Eric B. Fokam for help with permits and logistics; and the staff of the Mount Cameroon National Park and the Bokwango and Bakingili communities for their support. This study was performed with all the required authorisations of the Republic of Cameroon Ministries for Forestry and Wildlife and for Scientific Research and Innovation.

Our research was funded by the Czech Science Foundation (20-16499S and 18-10781S) and Charles University (UNCE204069).

## Author contribution

SPS and RT conceived the idea, RT and ŠJ designed the study, ŠJ, RT, YK, and JEJM sampled the data, YK, JEJM, and RT supervised processing of the video recordings, JH identified the floral visitors, SPS, YK, JEJM, and RT assessed the visitor behaviour, SPS analysed the data, SPS, RT, and ŠJ interpreted the results and wrote the manuscript draft, all authors commented the manuscript and approved its submission.

## Data availability

Data is available through Zenodo (*DOI will be provided upon acceptance*).

