## Supplementary material (tables 1-4, figure 1) for "Cheaters among pollinators: Floral traits drive spatiotemporal variation in nectar robbing and thieving in Afrotropical rainforests" (from another version of this preprint: version 1, 2022-05-16)

### *New Phytologist* Supporting Information

Article acceptance date: Click here to enter a date.

The following Supporting Information is available for this article:

**Table S1** An overview of the sites of the studied flower-visitor communities on Mount Cameroon.

**Table S2** Floral traits used in multivariate analyses with details of the categories used and further explanations where applicable.

**Table S3** A summary table on relationships among cheating behaviour, environmental variables, and spatiotemporal distribution of floral traits on Mount Cameroon as analysed by the RLQ analysis

**Table S4** Summaries of separate forward-selection RDAs of preferences of nectar robbers and thieves on Mount Cameroon.

**Fig. S1** Overview of the RLQ analysis used for analysing spatiotemporal associations of floral traits and visitor behaviour on Mount Cameroon

**Table S1** . An overview of the sites of the studied flower-visitor communities on Mount Cameroon.

| **Site** | **Elevation (a.s.l.)** | **Latitude and Longitude** | **Forest structure** | **Sampling year** |
| --- | --- | --- | --- | --- |
| Mann’s Spring | 2,200m | N 04.1428°, E 09.0630° | Montane primary forest just below the timberline | 2017 |
| Crater Lake | 1,450m | N 04.1443°, E 09.0709° | Submontane primary rainforest, lower tree density due to natural disturbances by elephants | 2017 |
| PlanteCam Camp | 1,100m | N 04.1175°, E 09.0717° | Upland primary rainforest, lower tree density due to natural disturbances by elephants | 2018 |
| Drink Gari Camp | 650m | N 04.1022°, E 09.1225° | Lowland primary rainforest, closed high canopy and scarce understory layer | 2018 |

**Table S2** Floral traits used in multivariate analyses with details of the categories used and further explanations where applicable.

| **Trait** | **Category or unit of measurement** | **Additional notes** |
| --- | --- | --- |
| *Morphological* | |  |
| Shape | Bell /Bowl / Dish / Funnel /Gullet / Labiate / Open / Papilionaceous / Salverform / Stellate / Trumpet / Tube / Urceolate | Following definitions from by Faegri and van der Pijl (1979), Ramirez 2003 and Ollerton and Watts 2002) |
| Symmetry | Actinomorphic / Zygomorphic |  |
| Orientation | Horizontal / Pendant / Upright / All | Species whose flowers were found to be orientated in all three directions were categorised separately into “All”. |
| Odour strength | None / Weak / Strong | Categorised by observer, hence kept simple |
| Brightness | Drab / Vivid |  |
| Colour | Brown / Green / Orange / Pink / Purple / Red / White / Yellow | Categorised by visual inspection |
| Nectar guides | Present / Absent | Categorised by visual inspection (in visible and UV spectra) |
| Size | Total corolla size (cm) | Zygomorphic flowers were measured along the horizontal and vertical axis |
| Tube width | Width of nectar tube (cm) | Measured at the centre of the nectar tube |
| Tube length | Length of nectar tube (cm) |  |
| *Nectar* | |  |
| Volume | (μL) | Measured as 24-hour production in flowers in-situ, as described in Janeček et al., 2021. |

**Table S3** A summary table on relationships among cheating behaviour, environmental variables, and spatiotemporal distribution of floral traits on Mount Cameroon as analysed by the RLQ analysis (see Fig. S4 for the conceptual details). The three matrices used were R (environmental variables and visitation frequencies for robbers, thieves, and pollinators per site), L (plant species composition per site), and Q (floral traits per plant species). (A) Summary of the separate analyses of R, L, and Q matrices: Eigenvalues and explained variation for axis 1 and axis 2, and the total variation of each matrix. (B) Summary of the RLQ analysis: Eigenvalues and percentages of total co-structure between R and Q represented by the two RLQ axes, covariance and correlation between the sites constrained by environment and visitor behaviours in R and plant species constrained by the traits in Q. (C) Projected inertia from individual analyses: Eigenvalues for matrices R and Q on the first two RLQ axes and the amount of projected inertia from the individual analyses preserved by the RLQ analysis.

| (A) **Summary from individual analyses of R, L, and Q** | | | | | |
| --- | --- | --- | --- | --- | --- |
|  | | | Axis 1 | Axis 2 | Total variation |
| **R** Environmental variables and visitation frequencies per site (*Hill-Smith Analysis)* | | | | | |
| Eigenvalues | | | 2.000 | 1.686 | 5.000 |
| Explained variation (%) | | | 40.17% | 33.73% | 73.90% |
| **L** Plant species composition per site (*Correspondence Analysis)* | | | | | |
| Eigenvalues | | | 0.86 | 0.77 | 4.25 |
| Explained variation (%) | | | 20.20% | 18.12% | 38.4% |
| **Q** Floral traits per plant species (*Hill-Smith Analysis)* | | | | | |
| Eigenvalues | | | 3.062 | 2.127 | 25 |
| Explained variation (%) | | | 12.24% | 8.50% | 20.75% |
| (B) **Summary of results from RLQ analysis** | | | | | |
|  | Axis 1 | Axis 2 |  | | |
| Eigenvalues | 0.427 | 0.147 |  |  |  |
| Explained variation in co-structure of R and Q (%) | 60.32% | 20.80% |  |  |  |
| Covariance | 0.65 | 0.38 |  |  |  |
| Correlation | 0.39 | 0.21 |  |  |  |
| (C) **Variation preserved from individual analyses for each RLQ axis** | | | | | |
|  | Eigenvalues | | Amount of projected variation | | |
|  | Axis 1 | Axis 2 |  |  |  |
| **R** Environment and behaviour | 1.647 | 1.925 | 96.67% | | |
| **Q** Floral traits | 1.677 | 1.587 | 62.92% | | |

**Table S4** Summaries of separate forward-selection RDAs of preferences of nectar robbers and thieves on Mount Cameroon. For each RDA, the table contains adjusted explained variance (R2adj), and F, and P-values (analysed by Monte Carlo permutations) for the whole model and for the floral traits with significant effects. ** p < 0.01, * p < 0.05.

|  | **Model** | | | **Floral traits** | | |
| --- | --- | --- | --- | --- | --- | --- |
|  | ***R^2^_adj_*** | ***F*** | ***p-value*** |  | ***F*** | ***p-value*** |
| Robbing | 0.18 | 3.22 | 0.001** | Tube length | 13.58 | 0.002** |
|  |  |  |  | Shape | 2.07 | 0.012** |
|  |  |  |  | Tube width | 3.47 | 0.036* |
| Thieving | 0.12 | 2.30 | 0.002** | Brightness | 3.20 | 0.008** |
|  |  |  |  | Shape | 1.97 | 0.014** |
|  |  |  |  | Tube length | 2.92 | 0.042* |

**Fig. S1** Overview of the RLQ analysis used for analysing spatiotemporal associations of floral traits and visitor behaviour on Mount Cameroon, modified from Dolédec et al. (1996). The analysis aims to reveal the co-structure between the two matrices R (environmental variables and behaviour frequencies) and Q (floral traits) through the central matrix L (species composition) acting as a link. Ordinations for matrices R and Q were independently conducted using Hill-Smith Analysis, while matrix L was subjected to Correspondence Analysis.

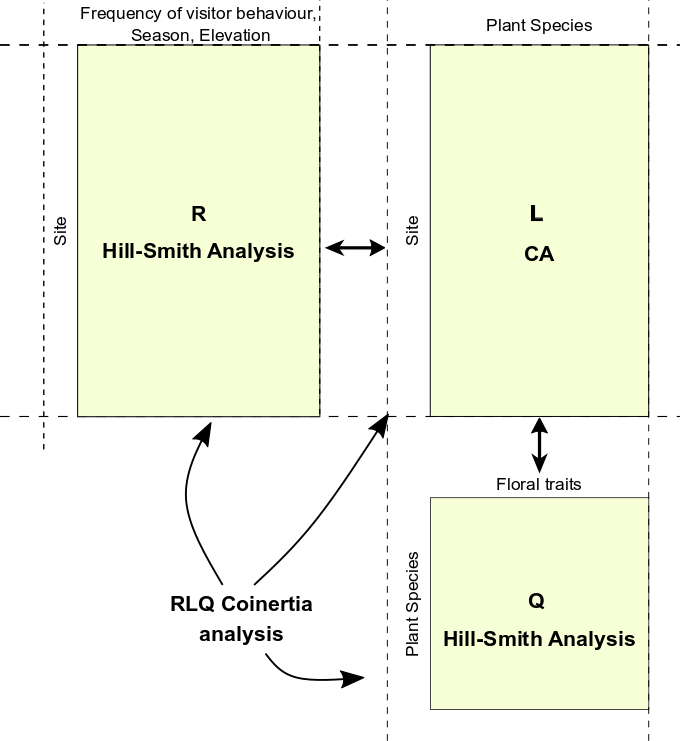
